# Peripheral tissues of deep-sea mussels exhibit autonomous circadian timing via an atypical mechanism

**DOI:** 10.1101/2025.10.08.681125

**Authors:** Audrey M. Mat, Federico Scaramuzza, Christophe Klopp, Marjolaine Matabos, Kristin Tessmar-Raible

## Abstract

While biological rhythms are crucial to life, the deep sea has long been considered an arrhythmic exception. However, at hydrothermal vents - devoid of diel cues yet shaped by tides - the mussel *Bathymodiolus azoricus* shows both tidal and, unexpectedly, circadian rhythms at −1700 m. Whether endogenous clock(s) drive these cycles remained unanswered. Here, we report endogenous circadian rhythms in *B. azoricus* cell cultures under constant conditions: isolated cells displayed a circadian oscillator despite tidal-dominant rhythms in situ. Reporter assays using genomic regions upstream of the mussel’s *per* gene and containing E-box motifs indicate that a functional transcription–translation feedback loop (TTFL) underpins circadian timing even in the deep sea. In contrast to conventional models, however, *BazPeriod* lacks autonomous repressive activity but modulates *BazCry2*. As *BazPeriod* itself oscillates tidally, it may explain how a single endogenous clock yields both tidal and diel rhythms. The work also spotlights the highly time-sensitive biology of vastly unexplored deep-sea biology.

## Introduction

Ocean ecosystems harbour remarkable biology, play a vital role in planetary health, and the ecosystem services they provide are crucial to humanity (*1*). Yet, the fundamental temporal biological principles that govern these ecosystems are complex and remain poorly understood. Timing is critical, as optimizing any given process to the right time can make the difference between life and death, especially at environments where environmental changes can be extreme. Thus, critical biological processes are often not merely direct responses to environment cues, but are driven by endogenous oscillators — commonly known as biological clocks — that enable organisms to anticipate predictable, regular, environmental changes (*2*, *3*).

The ocean is predominantly the deep ocean, which accounts for more than 90% of its total volume (*4*, *5*). It is defined as waters deeper than ∼200 m, where light is insufficient to support net photosynthetic production. Conventional wisdom initially propagated the view that these were cold, dark environments, with very few environmental changes, i.e. arrhythmic. However, it is increasingly recognized that the deep sea is a mosaic of ecosystems composed of numerous, diverse habitats and communities distributed heterogeneously (*6*), including hydrothermal vents. One of the habitat-defining key species living on hydrothermal vents is the deep-sea mussel *Bathymodiolus azoricus,* which inhabits vents along the Mid-Atlantic Ridge, at depths of 840 to 3350 meters (**Fig.1A**). There, pressures range from 84 to 335 bar (sea level: 1 bar), and temperatures range from 4°C in ambient seawater to 350°C in black smoker fluids. Mussel assemblages preferentially occupy cooler waters next to the vent fluids (4-9°C; *7*). Due to oceanic tidal pressure and modulation of local currents by ocean tides (*8*, *9*), the mussels’ environment alternates between the influence of anoxic, hypercapnic, acidic, metal-rich vent fluid and oxygenated surrounding seawater, creating a temporally highly dynamic, but also predictable, complex chemical landscape switching between life-enabling and harsh, toxic conditions (**Fig.1B**). Consistently, tidal environmental signals were recorded in temperature and pressure data from permanently deposited instruments in the field (*8*, *10*).

**Figure 1.**
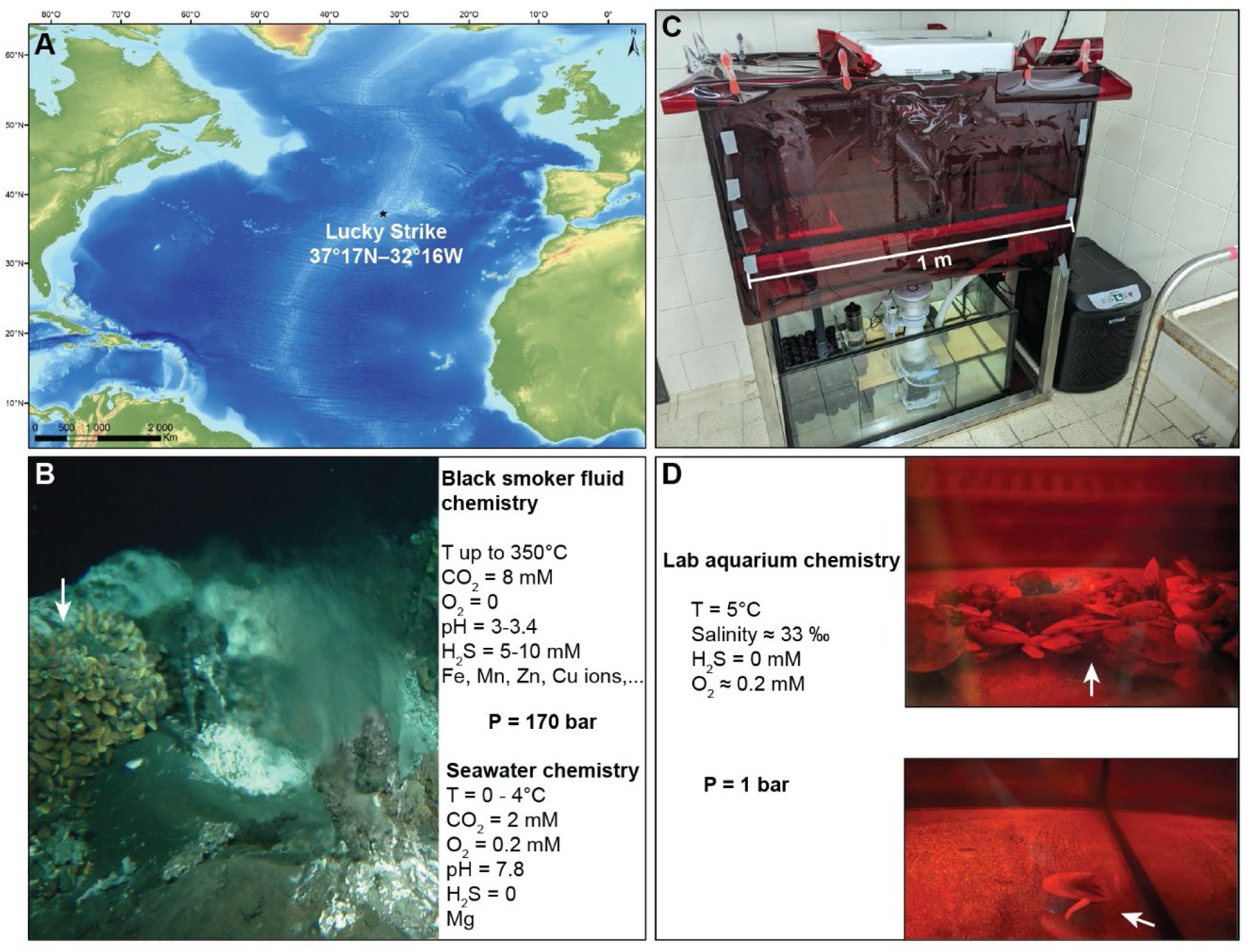
Sampling site of *Bathymodiolus azoricus* and comparison with the lab environment. **A.** The Lucky Strike vent field is located on the Mid-Atlantic Ridge at 1700 m depth, south of the Azores Triple Junction (*46*). **B.** Representative image of the environment, and conditions (*7*). Temperatures range from 4°C in ambient seawater, to 350°C in pure black smoker fluids. Mussels (arrows) inhabit areas around vents between 4 and 9°C. Photo @Ifremer. **C,D.** Representative images of aquarium set-up wrapped in red filters, and conditions. Mussels (arrows) moved regularly in the aquarium, on the walls or at the bottom.

Thus, environmental cycles are important for the local mussel communities. In line with this, recent work showed that on the deep seafloor, at −1688 m depth on the Lucky Strike vent field on the Mid-Atlantic Ridge (**Fig.1A**), the dominant rhythm of behaviour and at molecular transcriptomic level in *B. azoricus* is tidal (12.4 h), yet, strikingly, a significant circadian signal also emerges despite the absence of any clear diel cycle at vents (*10*). These findings matched well with previous work on shell growth and community dynamics in the Pacific that also suggested tidal cycles (*7*, *11*, *12*), and is further supported by data showing circatidal rhythms in vent shrimps *Rimicaris leurokolos* in constant conditions (*13*). However, it also results in questions on how these rhythms would change under different conditions. In fact, *B. azoricus* sampled at the nearby Menez Gwen vent field (−800m) and exposed to a 12:12 Light:Dark (L:D) cycle under controlled lab conditions exhibited mostly a diel cycle, with a persisting tidal cycle in RNAseq data (*10*). The presence of a weak circadian signal in the deep-sea, where there is no known daily cycle, and of a circatidal signal in the lab, where there was no tidal signal, suggested that these rhythms could be driven by one or two endogenous clock(s). Of note: while acknowledging that formal entrainment by tidal cues has not yet been demonstrated for *B. azoricus*, we use the term “circatidal” to describe the observed ∼12.4 h free-running rhythm as the mussels originate from a clearly tidal habitat.

At the molecular level, the core of the conventional circadian clockwork in animals relies on transcription-translational feedback loops (TTFL), including *clock* (*clk*), *bmal* (aka *cycle* in *Drosophila*), *period* (*per*), and in a species-dependent manner, *timeless* (*tim*), and different *cryptochromes* (cry; *3*, *14*, *15*). The transcript levels of several of the genes oscillate, as a relevant feature of the oscillatory mechanism (*3*). With the exception of echinoderms and hemichordates, where *per* genes have been lost (*16*), *per* transcripts cycle in all animals where they have been analysed, such as mice (*17*), flies (*18*), butterflies (*14*), crustaceans (*19*), and marine annelids (*15*). *Period* transcription is activated via specific regulatory enhancer sequences, called E-boxes, that are bound by the transcriptional activators CLOCK and BMAL (*20*–*22*). The resulting PER proteins repress CLOCK/BMAL’s action, being the negative loop of the core TTFL. *Tim* in *Drosophila* and mammalian *cry1, cry2* are also activated by CLK/BMAL via E-boxes, and form a heterodimer with PER in the negative part of the loop (*3*). While, of course, the global molecular circuitry of the circadian clock also involves a complex network of post-transcriptional, post-translational, and epigenetic processes (*3*, *23*), the above mentioned mechanism is at the core of the TTFL in flies, mice and humans (*3*).

Based on the previous identification of core circadian clock gene orthologs in *B. azoricus* (*10*), we wondered if these may be part of a TTFL core loop mechanism. Considering the presence of molecular circadian clocks in various cells and tissues (*24*–*26*), we also wondered whether we could find evidence for circatidal and/or circadian clock(s) in the tissues of *B. azoricus*. The establishment of *B. azoricus* primary cell cultures for chronobiological analyses and subsequent long-term (> one year) maintenance allowed us to approach these questions on the tissue level, and to dig into the mechanism of the clock(s) in deep-sea mussels, that do not possess a centralized brain (*27*).

## Material and Methods

### Sampling procedure and animal care

All sampling and animal work were performed according to the legal European and national guideline. The samplings comply with the Nagoya regulation. The Lucky strike vent field is part of the Exclusive Economic Zone of Portugal, and the Portuguese Authorities issued Internationally Recognized Certificate of Conformity (IRCC) ADENDA 22/2022/DRCTD for the Momarsat22 cruise, and IRCC ADENDA CCIR-RAA/2023/35 for the Momarsat23 cruise. The maintenance of the mussels does not require additional animal experimental approval as these are non-cephalopod invertebrates.

In 2022, the mussels were collected at 1694 m depth at the “Tour Eiffel” edifice, using the *Nautile* submarine during the Momarsat22 cruise (*28*) on June 22 (dive 2039-12). In 2023, the mussels were collected at 1700 m depth at the “Montsegur” edifice, using the Remotely Operated Vehicle (ROV) *Victor6000* during the Momarsat23 cruise (*29*), on July 24^th^ (dive 846-8). Mussels were kept onboard until the end of the cruise (6 and 7 days in 2022 and 2023, respectively) in darkness at 8°C, and water was replaced every day. They were then shipped to Vienna by plane within 24h (Flying Shark Lda, Portugal). In 2022, the survival rate of mussels upon transport was 86 %. In 2023, the survival rate of mussels after transfer was 70 %.

In 2022, mussels were housed in a 36-L aquarium in a cold room at 4.3 ± 1.0°C (mean ± SD) under a 9:15 Light:Dark cycle (white LED, 68 Lux), with a salinity of 35.3 ‰. Water was changed once or twice a day manually. In 2023, mussels were housed in a 150-L aquarium with a filtering system, and kept at 5.5 ± 0.5°C with a salinity of 33.1 ± 1.0 ‰. The aquarium was sheltered from the 16:8 L:D cycle of the room by 026 – bright red filters (Cotech Sensitising Ltd, UK; **Fig.1C, D**). After 1 month, mussels were fed with grinded fish flakes (TetraMin, Tetra GmbH, Germany).

### Cell cultures

Though rare prior successes cultured cells from shallower species (*30*, *31*) or, at comparable depths (*32*), only for a few days, we wanted to develop tissue cultures for *B. azoricus*. The dissociation protocol is a modified version of the cell dissociation of *Platynereis* whole larvae protocol (*33*). All steps until the cells’ distribution in 6-well plates were performed on ice, unless stated otherwise. Mussels were dissected, and the pieces of tissue (20-70 mg) were directly transferred into individual eppendorfs with 1 mL filtered seawater and 2% penicillin / streptomycin (ref. P4333, Sigma-Aldrich, USA). The samples were allowed to sink to the bottom; then, supernatant was removed and replaced again by 1 mL filtered seawater and 2% penicillin / streptomycin. Supernatant was removed again and replaced by 400 µL filtered natural seawater with 0.15 µg/µL pronase (Sigma-Aldrich, USA) and 10 µg/µL sodium thioglycolate (ref. T0632-25g, Sigma-Aldrich, USA). Samples were incubated at room temperature for 10 minutes, then rinsed twice with 1 mL PBS (phosphate buffered saline, ref. P5493, Sigma-Aldrich, USA). Supernatant was then removed, and samples were incubated in 400 µL with PBS with 150 µg/mL of liberase (ref. 05401020001, Roche, Germany). Again, samples were rinsed twice with 1 mL of PBS. The next step was performed under a laminar flow hood: samples were dissociated in 1 mL PBS with 3 µL 10% bovine serum albumin (ref. A3912, Sigma-Aldrich, USA). They were first pipetted up and down for 5-10 minutes with a P1000 then a P200. None of the samples ever fully dissociated. The supernatant was then passed through a 70 µm Flowmi cell strainers (ref. 136800070, SP Bel-Art, USA) and seeded in a well of a 6-well plate (CC7682-7506, Cyto One, Germany). Four mL of medium, prepared according to (*34*), were then added to each well.

Cells were cultured in tissue culture treated 6-well plates, then transferred to T-25 flasks (CC7682-4325, Cyto One, Germany). All cultures were incubated under constant darkness. Several cultures were imaged using a Zeiss Observer Z1 inverted microscope (Zeiss, Germany). Besides using non-coated plates or flasks, three different coatings have been tested on foot cultures: fibronectin (Yo Proteins, 663, Sweden) diluted 1:100 in PBS, gelatin (G1890, Sigma-Aldrich, USA) 0.18% in PBS, and Cultrex Ultimatrix (BME001, Biotechne, USA). Cultures did not adhere with neither gelatin, nor fibronectin, nor Cultrex Ultimatrix coating. These coatings actually all quickly favoured the growth of mould (n = 2 cultures tested for each coating).

The identity of the cell cultures has been evaluated by PCR for foot, mantle, and muscle tissues. The PCR targeted the ribosomal protein L15 (RPL15) with primers as in (*35;* Forward: TATGGTAAACCTAAGACACAAGGAGT; Reverse: TGGAATGGATCAATCAAAATGATTTC).

Foot cell cultures have been confirmed after 83 and 184 days, the latter being further confirmed by Sanger sequencing. Similarly, a foot culture from the 2023 batch has been confirmed by RNAseq after 165 days. Two methods were tested to extract total RNA: the Direct-Zol RNA miniprep kit (Zymo Research, USA) using RNAzol (Sigma-Aldrich, USA), and a phenol:chloroform protocol with Extract-all (Eurobio, France) according to manufacturer instruction. Out of the 2 RNA extraction methods tested, the phenol:chloroform protocol proved to be the most efficient one, allowing to recover 2 times more RNA from cell cultures. One should note that RNA concentrations could only be measured from a full culture from a 6-well-plate well or T25 flask, and even so, detected RNA was around the detection limits of the instrument (2 ng/µL; Nanodrop 2000c, Thermo Scientific, USA). As a comparison, both methods were also tested on tissues: on average, the phenol:chloroform allows to extract 7 times more RNA than the RNAzol kit (19.7 ± 3.8 ng.µL^-1^.mg_tissue_^-1^ versus 2.9 ± 1.6 ng.µL^-1^.mg ^-1^, respectively). We also tested different volumes of Extract-all: 200, 300, 400, 600, and 800 µL on a smaller cell culture. RNA concentration was below the detection limit of the instrument; but PCR bands appeared similar on a 2% agarose gel. Phenol:chloroform therefore appeared as the method of choice for low input RNA recovery. Volumes of 400 µL of Extract-All or higher were easier to manipulate.

The optimal processing of the cultures appeared to be as follow: changing medium or transferring the cells at day 3, day 6, then taking care of the cultures every 5-7 days. Adherence of foot cells to the well or flask requires several days. Therefore, changing the medium every day at the beginning of the cell culture is counter-productive and leads to the loss of cells. Also, it should be noted that even on the densest cultures, no pellet was ever visible after centrifuging the supernatant. One therefore works blindly and need to limit the processing of the cultures. Once cells attach, handling the cultures becomes easier in terms of medium change. For longer term maintenance, once the culture appears stable, medium can be changed every 2, 3, or 4 weeks without noticing any visual, qualitative, difference.

Gills tissues are also promising tissues for future trials: cells would either adhere or stay in suspension, with homogenous round cell type of similar size and great densities initially close to 70-80 % confluence. However, all cultures that were started became contaminated within the next 12 days (n = 18, 2023 mussel batch) by unicellular organism swimming within the cultures. Microscope observations showed the presence of 2 flagellas, and kraken analyses on sequenced cultures furthered pointed towards Bicosoecida, and specifically *Cafeteria roenbergensis*, a protist that has been identified throughout the water column, including at hydrothermal vents (*36*, *37*). A co-culture of *B. azoricus* gill cells and *C. roenbergensis* has been maintained in the lab for > 2 years at 4°C.

### Temporal experiment

Animals were acclimated for four days after their arrival in the lab, and subsequently 5 mussels were dissected on ice. Foot tissues were cultivated as described above, at 4°C. This is day 0 for the cell cultures. Cultures were transferred to T-25 flasks on day 3, and medium was changed on day 6, 12, and 19 (**Fig.2**). On day 24, for each culture, cells from one T25-flask were resuspended, counted with a Neubauer chamber, and seeded in 24-wells plates (CC7682-7524, Cyto One, Germany; one plate par time point) with 1 mL of fresh medium at a density of ∼45 000 cells/well. Each medium change also represented a light pulse and a temperature shock, as cells were handled at room temperature in a laminar flow hood. Cells were incubated under constant conditions (4°C, constant darkness) for 24 h, then sampled every 4 h for 48 h. For each sampling point, cells were re-suspended by pipetting up and down and using the tip as a scrapper. They were then transferred to an Eppendorf and centrifuged at 400 g for 4 min. Supernatant was then removed, cells were flash frozen in liquid nitrogen, and stored at −70°C until further processing. Total RNA extraction was later performed using Extract-all (Eurobio, France) according to manufacturer instruction, using 400 µL of reagent per sample. RNA concentrations were below the detection limit for Nanodrop or Femto Analyzer measurements, hence quality controls were performed bioinformatically after sequencing. Libraries were prepared with the SMART-Seq3 protocol (*38*). Samples were multiplexed and pair-end sequenced on a quarter of a NovaSeq S4 lane (Illimina Inc, USA) at the Vienna Biocenter Core Facilities GmbH, Austria.

**Figure 2.**
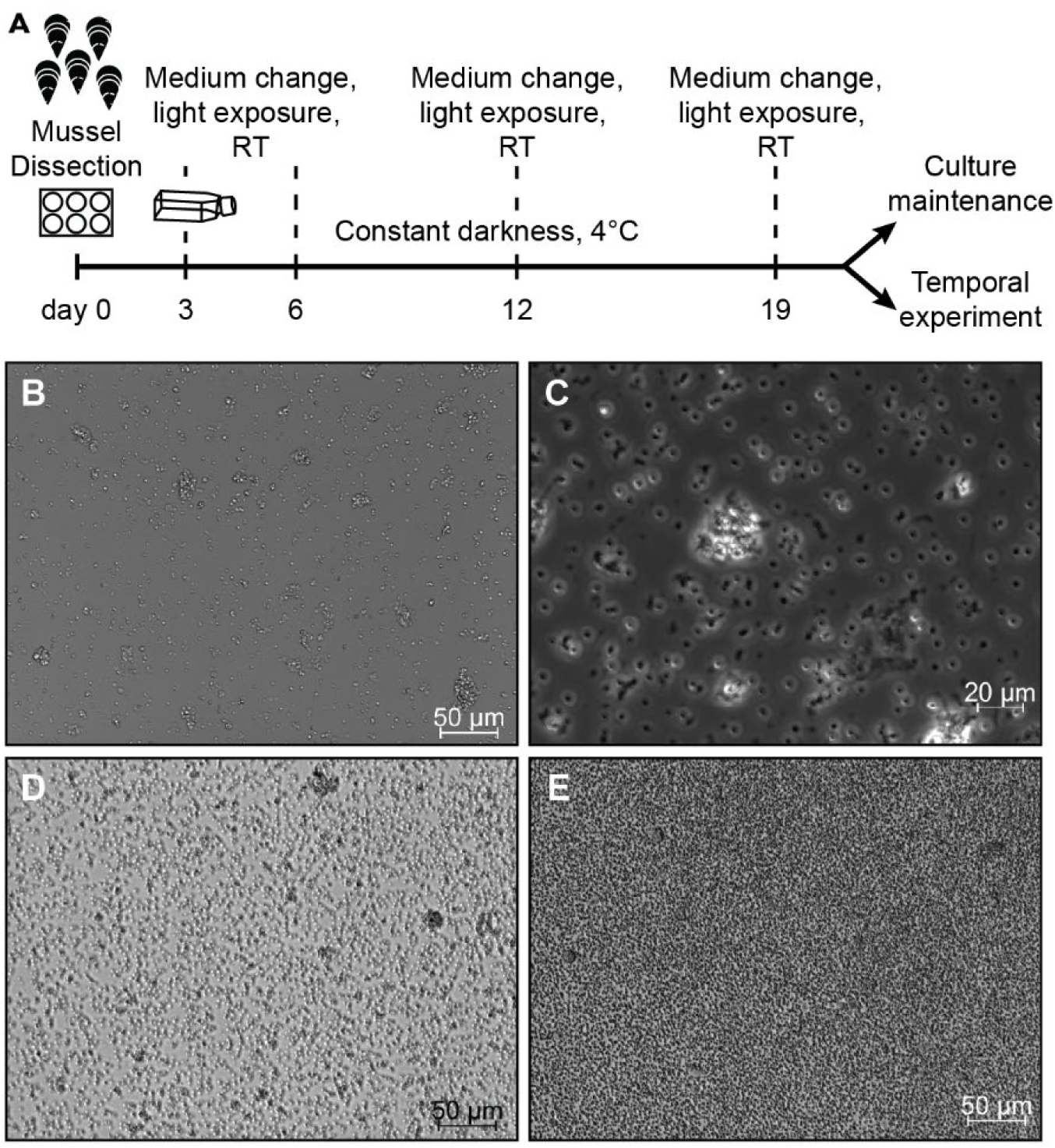
Cell cultures from *Bathymodiolus azoricus* tissues. **A.** Scheme of the protocol used to establish primary cell cultures from *B. azoricus*. RT: Room Temperature. **B, C.** Cell cultures from foot tissues, representative of the ones used in the subsequent experiment, imaged after 164 days. **D, E.** Cell cultures from mantle tissues, imaged after 15 days.

The sequencing of the 65 samples (13 time points x 5 mussels) provided 15.9 ± 2.8 million reads per sample (mean ± SD) before filtering. Reads were aligned on the reference transcriptome of *B. azoricus* (NCBI accession: HBDL01000000.1) using bwa mem (version 0.7.17, default parameters). The alignments were sorted, compressed and indexed using samtools sort and index (version 1.14, default parameters). The alignment rates were comprised between 36 and 81 %, with an average at 59 ± 11 % (mean ± SD). Quantification was then performed with samtools idxstats (version 1.14, default parameters). We performed an additional filtering after sequencing to evaluate the quality of the input RNA for each sample. CIGAR lines were extracted from the BAM alignment files; they describe how a sequenced read aligns to the reference. Only CIGAR lines of primary alignments were kept. Aligned fractions were calculated by dividing the match and mismatch length (sum of M sections) by the read length, 150 nucleotides. Then, the number of alignments were calculated for each given matched fraction. Based on the calculated profiles for each sample, we filtered out samples with low quality reads because if *B. azoricus* RNA is present in the sample, most of the alignments should be long (see Results). After filtering, the alignment rates ranged between 47 and 81 %, with an average at 64 ± 9 % (mean ± SD). Then, transcripts where the count was 0 for 35 or more of the samples were filtered out, as performed in (*10*). For comparison again (*10*), the data were normalised using DESeq2 (*39*; v 1.38.3) without using a zero-centered normal prior. The analyses were then run on the median counts for each time point using RAIN (*40*; v 1.32.0). To validate the results, a bootstrap approach was used: shuffling the columns of the data, running the RAIN analysis on the shuffled columns, and actually bootstrapping that procedure 10,000 times. The results were then compared with the original data. The analyses were run in R (*41*; v 4.2.3). Gene ontology enrichment analysis was run using the topGO package (*42*; v 2.50.0)

### Cloning of putative core circadian clock genes

Mussel RNA was extracted from mantle, gill, or foot tissues using Extract-all reagent (Eurobio, France) using 1 ml·30 mg^−1^ tissue. cDNA was produced using the LunaScript RT SuperMix Kit (New England Biolabs, USA). Full-length coding sequences of *B. azoricus clock* (*BazClk*), *bmal* (*BazBmal*), *period* (*BazPer*), *timeless* (*BazTim*), light-receptive *cryptochrome 1* (*BazCry1b*), and transcription-repressor *cryptochrome 2* (*BazCry2*) were then amplified by PCR from cDNA using the primers given in **Table S1** with a Phusion® High-Fidelity DNA Polymerase (New England Biolabs, USA). The gene *BazClock* appeared 57 nucleotides shorter compared to the previously published sequence (*10*). Similarly, *BazCry1b* was 6 nucleotides shorter. And for *BazBmal*, 56 nucleotides that were previously not identified within the first PAS domain have now been characterized. The sequences were directionally cloned into the pAC5.1_V5-HisA plasmid using the restriction enzymes given in **Table S1**. Competent *Escherichia coli* (NEB® 5-alpha Competent *E. coli*, New England Biolabs, USA) where transformed with the respective plasmids, and grown overnight in LB medium with 1:1000 (vol:vol) ampicillin. The plasmids were recovered with mini- or maxiprep kits (QIAGEN Plasmid Kits, Qiagen, Germany) and controlled by Sanger sequencing (Microsynth AG, Switzerland).

### Identification and cloning of putative enhancer constructs

To identify a potential enhancer construct in *B. azoricus*, we compared the genomes available for the deep-sea mussels *Gigantidas platifrons* (GenBank accession GCA_002080005.1), *Bathymodiolus brooksi* (Genbank accession GCA_963680875.1), and *Bathymodiolus septemdierum* (Genbank accession GCA_963383655.1). We first blasted the sequence of *BazPer* (GenBank MN611455) against the 3 different genomes to identify the contigs and the positions of the coding sequence of the respective *period* genes. We then extracted the nucleotide sequence 100 kb before the first coding exon for the 3 species. We compared the 100 kb sequences of *G. platifrons* and *B. brooksi* using Mulan (https://mulan.dcode.org/; *43*).

In this window, there are 9688 Ns in the sequence of *G. platifrons*, and none in the sequence of *B. brooksi*. *Bathymodiolus azoricus* gDNA extraction was performed using 50 µL of 50 mM NaOH, incubated at 95°C for 20 minutes, followed by the addition of 5 µL of 1 M Tris-HCl (pH 7.5). The potential enhancers were amplified from gDNA by PCR using the primers given in **Table S1** with Phusion® High-Fidelity DNA Polymerase (New England Biolabs, USA). The sequences were blunted and ligated into the pJET1.2 vector (CloneJET PCR Cloning Kit, Thermo Scientific, USA). Besides the potential *B. azoricus* enhancer, we used the hsp70 core promoter from *Drosophila melanogaster* (kindly provided by the Stark lab). Three sequences were amplified by PCR with primers containing specific 10 to 15 nucleotides overlaps (**Table S2**) for further insertion into pGL3 basic (Promega, USA). Bands were controlled on a 2% agarose gel: specific PCR products were purified using either the Monarch PCR & DNA Cleanup kit (New England Biolabs, USA) or the Monarch® DNA Gel Extraction Kit (New England Biolabs, USA). The pGL3 vector was linearized with SmaI (New England Biolabs, USA). Three different constructs were prepared using directional cloning based on the homology sequences (In-Fusion Snap Assembly, Takara Bio Inc., Japan): 1) *Baz*Block108-*Dm-hsp70*::*Luciferase* using molar ratios of 2:5:1, 2) *Baz*Block118-*Dm-hsp70*::*Luciferase* using molar ratios of 2:5:1, and 3) *Baz*Block108-block118-*Dm-hsp70*::*Luciferase* with molar ratios of 2:2:1. Competent *E. coli* (NEB® 5-alpha Competent *E. coli*, New England Biolabs, USA) where transformed with the respective constructs using 5 µL of the In-Fusion reactions and grown overnight in LB medium with 1:1000 ampicillin. The constructs were recovered with mini- or maxiprep kits (QIAGEN Plasmid Kits, Qiagen, Germany) and controlled by Sanger sequencing (Microsynth AG, Switzerland). We used directed mutagenesis to modify the *Baz*Block118-*Dm-hsp70*::*Luciferase* construct. This construct contains 2 E-boxes like (CACGTT) and one E-box (CACGTG), and we mutated the different motifs to generate all the possible combination of the construct: one construct where the E-box-like 1, or the E-box, or the E-box-like 2 was mutated, one construct where both the E-box-like 1 and the E-box were mutated, one where the E-box-like 1 and 2 were mutated, one where the E-box and the E-box-like 2 were mutated. The E-box-like 1 CACGTT was mutated to GCGCCG, the E-box CACGTG was mutated to TGCGCG, and the E-box-like 2 was mutated to GGGACA. The mutated motifs were selected to match the melting temperature requirements of the primers (**Table S3**) used with the In-Fusion Snap Assembly (Takara Bio Inc., Japan).

### Luciferase assays

*Drosophila* Schneider 2 (S2) cells luciferase assays were performed as described previously (*44*, *45*). In brief, cells were cultivated in S2 medium with heat-inactivated FBS (Sigma-Aldrich, USA) and 1% ʟ-glutamine-penicillin-streptomycin (Sigma-Aldrich, USA) at 25°C. The day before transfection, 400 000 S2 cells were seeded in 12-well dishes (Cyto One, Germany) in 1 mL of medium. On the day of transfection, the different plasmids were transfected into S2 cells using the Effectene® Transfection Reagent (Qiagen, Germany). We transfected either a total of 500 ng DNA to test the constructs or 1000 ng DNA to test the potential repressors, using the pAC5.1_V5-HisA plasmid to adjust the reactions to the final amount. We applied the kit provider’s instructions, using 4 or 8 µL of enhancer and 10 or 20 µL of transfection reagent for 500 or 1000 ng of plasmid, respectively. While complex formation took place, we added 300 µL of medium to each well, and 600 µL of medium to the tubes containing the transfection complexes. The respective complexes were then transferred drop by drop the corresponding wells. Plates were incubated for approx. 44 hours at 25°C in constant darkness before reading.

The cells were lyzed and processed with the Dual-Luciferase® Reporter Assay System from Promega (USA). The *B. azoricus* luciferase signal was normalised using the actin5c::Renilla luciferase. Both light signals were read using a BioTek Synergy H1 microplate reader (Agilent, USA) with a 2s-delay and a 8s-reading protocol. For each assay, at least 3 independent transfections were performed. Within each transfection, each condition was tested in biological triplicates. For each biological replicate, luciferase readings were measured in three technical replicates. Significant differences were evaluated by a one-way ANOVA with post-hoc Holm-Šídák multiple comparisons; SD are provided at the transfection-level.

## Results

### Establishment of long-term mussels’ maintenance in the laboratory

*Bathymodiolus azoricus* mussels were sampled on the Lucky Strike vent field on the Mid-Atlantic Ridge at approx. −1700 m (**Fig.1A, B**). In order to develop molecular tools and assays for *B. azoricus* and to lower the impact on the natural environment, we aimed for pushing the boundaries of deep-sea mussel maintenance in the laboratory. Individuals were sampled and transferred to the lab. The first batch of mussels was kept alive in the lab for a maximum of 12 days. We then significantly improved the system: mussels collected in 2023 were housed in a 150-L aquarium with a filtering system and protected from white light by red filters. From the pool of animals that were alive after transfer, 40 % died within the next 2 weeks, while 15 % visually healthy animals were sampled for cell cultures or RNA extraction. After 31 days, there were 51% dead animals, and 17% had been sampled. From then on, mussels were fed with grinded fish flakes. The remaining mussels not only survived in the lab, but lived apparently healthy lives, indicated by the observations that they occasionally changed their place in the aquarium, and were regularly observed open, sometimes with their foot out (**Fig.1D**). Animals remained alive for a total of 13 months, significantly expanding our ability to study them.

### Establishment of *Bathymodiolus azoricus* cell cultures

In order to assess the presence of possible endogenous cellular molecular oscillators, we established cell cultures. We tested dissociated cells from foot, gills, mantle, mantle edge, digestive gland, and muscles, and incubation temperatures at 4°C, 8°C, and 25°C (**Fig.2A**). No visual difference could be observed between cell cultures at 4 or 8°C. At 25°C, however, cells were quickly overgrown by fungi and other contaminants.

Among the different cell cultures (n = 63 primary cultures), foot cells were the most viable. The cells were round shaped with a diameter of 3.8 ± 0.2 µm (mean ± SE) and consistently adhered to the bottom of the wells or flasks (**Fig.2B, C**). The others cultures had to be discarded within the first 2 months, because of cell death or contamination. Mantle cells divided substantially within the first days, but none of these cultures could be successfully maintained beyond 2 weeks (n = 9 cultures). Mantle and gill cells are round shaped with a diameter of 4.0 ± 0.2 µm (mean ± SE) for the mantle (**Fig.2D, E**). The *B. azoricus* cell culture identity was confirmed by PCR for foot, mantle, and muscle tissues, and in part by sequencing for foot tissues (**Fig.S1A**).

Foot cell cultures could be transferred to another well or a T25 flask within the first days and weeks. However, none of the transfer that was attempted after a prolonged period in the same container (>1.5 months) was successful. Because cells could not be transferred, the assessment of the cultures has most of the time been performed visually, carefully observing the cultures over time, rather than counting cells on a regular basis, which would have terminated the cultures rapidly. Indeed, cultures were actually mainly established in the first ∼15 days. Thereafter, if density did not change dramatically, the cells showed signs of division: areas that had been scrapped for attempted transfer were slowly recolonized in several cultures, over several weeks or months. Two cultures are still being maintained beyond 3 years (**Fig.S1B-E**).

### Circadian rhythms are detectable in the transcriptome of foot cell cultures

In several species it has been shown that tissue cultures exhibit endogenous circadian clock oscillations (*24*, *47*–*49*). Cells in tissue cultures typically phase synchronize their circadian clocks after passaging or medium change (*50*). Mechanistically, a wide array of signalling pathways were shown to elicit circadian gene expression in cell cultures, including glucocorticoids (*51*), cAMP, protein kinase C, and Ca^2+^ (*52*). These molecules induce *per1* and *per2* transcription (*50*), in a similar way to light pulses in rats maintained under constant darkness (*53*), causing phase synchronization between the cells. Furthermore, circadian rhythms in tissue cultures can also be synchronized via environmental factors such as light (*49*, *54*) or temperature (*55*). Given the relatively general occurrence of this phenomenon, we reasoned that if endogenous circadian clocks are present in the cells and tissues of *B. azoricus,* they may also be synchronized by changing the medium and subsequently, we could be able to detect rhythmicity in their transcriptome. It should be noted that medium change is also an environmental shock for the cells (light and temperature). Whether similar rules also apply for potential circatidal clocks is unclear, as their mechanistic nature is not yet understood (*19*, *56*, *57*). We sampled cells under constant conditions every 4hrs over 2 days (**Fig.3A**). Sampling duration and frequency was limited by the availability of the cell cultures.

**Figure 3.**
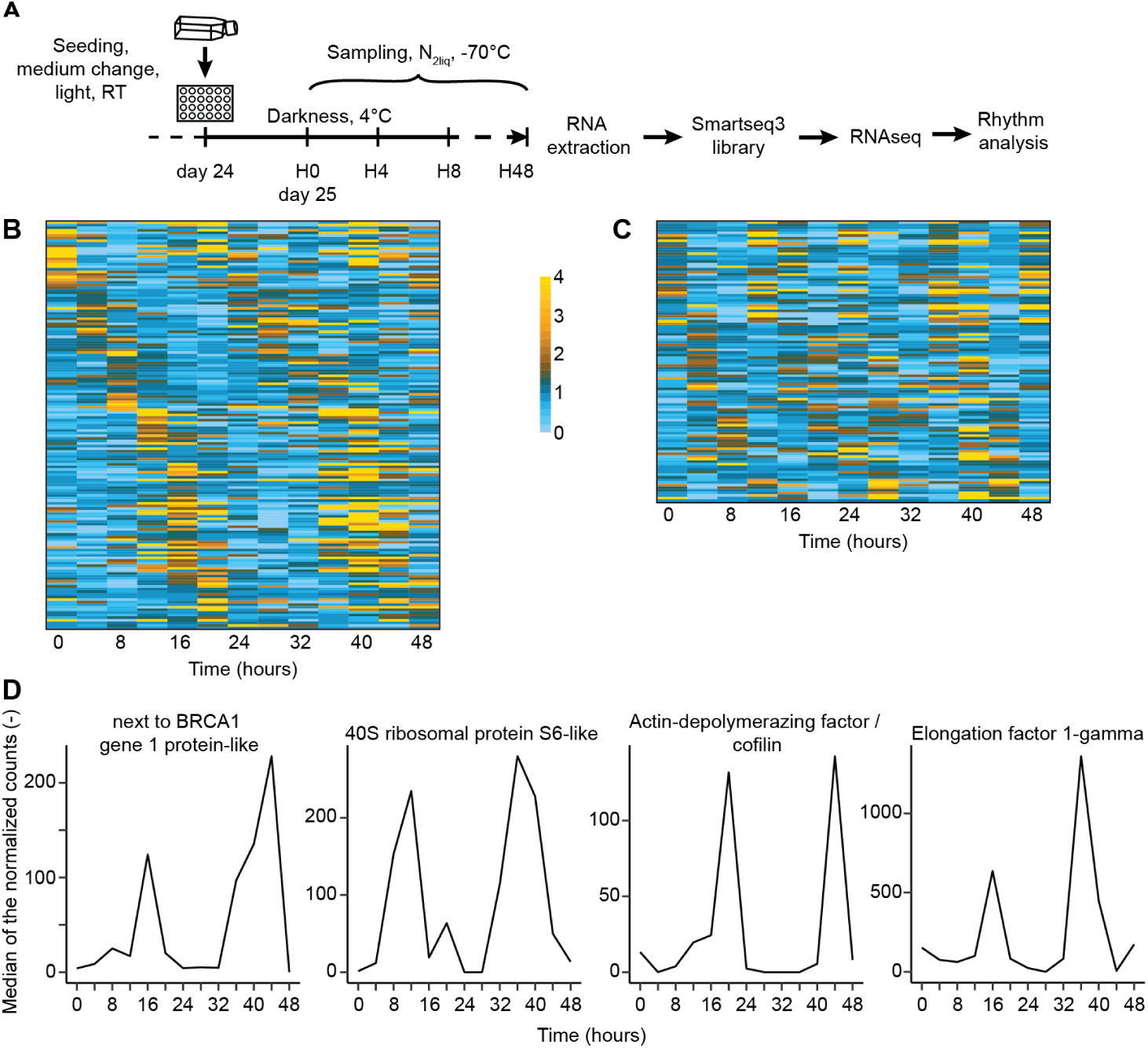
Identification of rhythmic transcripts in cell culture temporal experiment. **A.** Sampling procedure and processing of the data. RT: Room Temperature. **B.** Median-normalised expression of the 157 transcripts identified in the circadian range. **C.** Median-normalised expression of the 109 transcripts identified in the circatidal range. Transcripts are ordered by phase. Heatmap colours: median-normalised expression levels greater than 2-fold are shown with a yellow gradient; expression levels < 1-fold are shown with a light blue gradient. **D.** Plot of selected circadian transcripts identified using RAIN. Named according to best BLAST hit on protein level.

The sequencing of the 65 samples from the temporal experiment provided 12.8 ± 2.8 million reads per sample after read quality filtering (mean ± SD). Given the slow growth of the cells and the limited availability of animals, we worked with very low amounts of material. We therefore performed a quality control after sequencing based on the CIGAR lines of the BAM files (see Material & Methods and **Fig.S2**). The X axis represents the matched fraction from 0 to 100 %, which is calculated as the number of matching nucleotides divided by the total length of the read (150 nt). The Y axis represents the number of aligned reads for each matched fraction. Based on these profiles, the samples with less than 500k alignment counts at the 150 nt peak were removed from further analyses (**Fig.S2**). Then, filtering out transcripts for which ≥35 samples had 0 count yielded a shorter list of transcript: 5 616 transcripts out of 34 196 were kept, versus 34 163 and 34 141 out of 34 196 transcripts in the deep-sea and lab experiments in (*10*), respectively. RAIN analysis identified 157 transcripts exhibiting a circadian period in the cell culture temporal experiment (**Fig.3B, D, Table S4**). Among these 157 transcripts, 66 of them had previously been identified as rhythmic (tidal/circatidal; diel/circadian) in gill RNAseq data from *B. azoricus* from the deep seafloor or the lab (*10*). Forty of those 157 transcripts had previously been identified specifically as daily or circadian (*10*).

We also tested for transcript rhythmicity in the circatidal range using RAIN and identified 109 transcripts exhibiting a circatidal period (**Fig.3C, Table S5**), of which 41 had previously been identified as rhythmic and 26 as tidal or circatidal in *B. azoricus* gill RNAseq data (*10*).

To further validate our analyses, we tested whether our results differed significantly from random expectations by analysing how frequently these numbers could arise by chance in randomised datasets. We randomly shuffled the timepoints in the dataset, and re-ran the rhythmic analyses, bootstrapping this procedure 10 000 times. For the circadian analysis, the number of rhythmic transcripts in the actual dataset always significantly outperformed the shuffled data, meaning that we got more than random (>50 % for each test, **Table S6**). In our experiment, we detected more circadian transcripts over random in 70.3 % of the cases, more circadian transcripts that were already identified as rhythmic in the gill RNAseq analyses on the deep seafloor or in the lab (*10*) in 77.9 % of the cases, and more circadian transcripts that were already reported as circadian in the gill RNAseq dataset in 90.8 % of the cases. Also, none of the 10 000 shuffled datasets led to the identification of at least 157 circadian transcripts, with the same 66 transcripts that were previously identified as rhythmic (**Table S6**). This strongly supports the existence of circadian transcripts in the free-running cell culture temporal experiment, and therefore of a circadian clock in deep-sea mussels’ cells. The enrichment analysis of gene ontology (GO) terms associated with biological processes revealed 35 enriched terms with *p*-value < 0.01, highlighting macromolecules/peptides/amide biosynthesis, regulation of gene expression (translation and post-translation), structural organization (centriole, cytoskeleton), and autophagy (**Fig.3D**, **Table S7**).

For the circatidal transcripts, we found no evidence for significantly better performance of our experimental data over the random shuffle (**Table S6**). However, there are multiple reasons that may explain why we did not observe circatidal oscillation in our tissue culture experiment, including the question of tidal phase synchronization of tissue cultures.

Of note, the transcripts of putative core circadian clock genes, like most transcription factors, could not be detected in our data, likely due to the limitations of the material and, hence, low sequencing depth.

### A core TTFL likely underlies the *Bathymodiolus azoricus* cellular circadian clock

As the transcriptomic analyses from cell cultures strongly suggest the presence of an endogenous, cell-autonomous circadian clock, we next wondered if this clock may function according to the principle mechanisms of the well-established TTFL.

The full sequences of the previously predicted core canonical circadian clock genes (*10*) were successfully cloned from *B. azoricus* cDNA (see Methods). In order to test if the putative core circadian clock genes of *B. azoricus* may function in a realistic deep-sea mussel genomic context, we searched for potential circadian-relevant enhancer regions in *B. azoricus*. In the absence of a *B. azoricus* genome, we compared the genomes available for the deep-sea mussels *Gigantidas platifrons* (previously known as *Bathymodiolus platifrons*), *Bathymodiolus septemdierum*, and *Bathymodiolus brooksi*. Circadian enhancers are typically located upstream the *period* gene in mammals and flies, and E-box sequences, usually CACGTG, are required for robust RNA cycling of the targeted genes (*20*). We blasted *BazPer* against the 3 available deep-sea mussel genomes and analysed the sequence spanning 100 kb before the position of the first *per* coding exon. We then used the MULAN algorithm to detect potential functional regulatory elements. As the sequences we analysed showed a very high overall conservation between *B. brooksi* and *B. septemdierum*, we further focused on the comparison between *G. platifrons* and *B. brooksi,* which allowed better discrimination between conserved and variable sequences. Conserved regions between more distant phylogenetic species typically correlate with conserved functionality. We identified 2 ortholog blocks containing the E-box CACGTG: block 108 and block 118 (**Fig.4A)**. Block 108 is located 12 769 and 19 283 nt upstream of *per* in *G. platifrons* and *B. brooksi*, respectively. Block 118 is located 7 605 and 5 957 nt upstream of *per* in *G. platifrons* and *B. brooksi*, respectively. After aligning the 2 blocks between the 2 mussel’s species, we designed degenerate primers and cloned the corresponding block 108 and block 118 from *B. azoricus. Baz*Block 108 is 421 nt long, and contains 1 CACGTG E-box located 11 nt before the end of the block. *Baz*Block 118 is 638 nt long and contains 1 CACGTG E-box and two CACGTT E-box-like ranging over 86 nt, located on the 5’-end of the block (**Fig.S3**). The configuration is similar in *G. platifrons* and *B. brooksi,* but *G. platifrons* does not possess the second E-box-like motif in block 118 (**Fig.4A**). The conservation of the block is specific to the identified sequences. Low sequence similarities are shown as representative examples 450 nt upstream and downstream of block 118 (**Fig.4A**).

**Figure 4.**
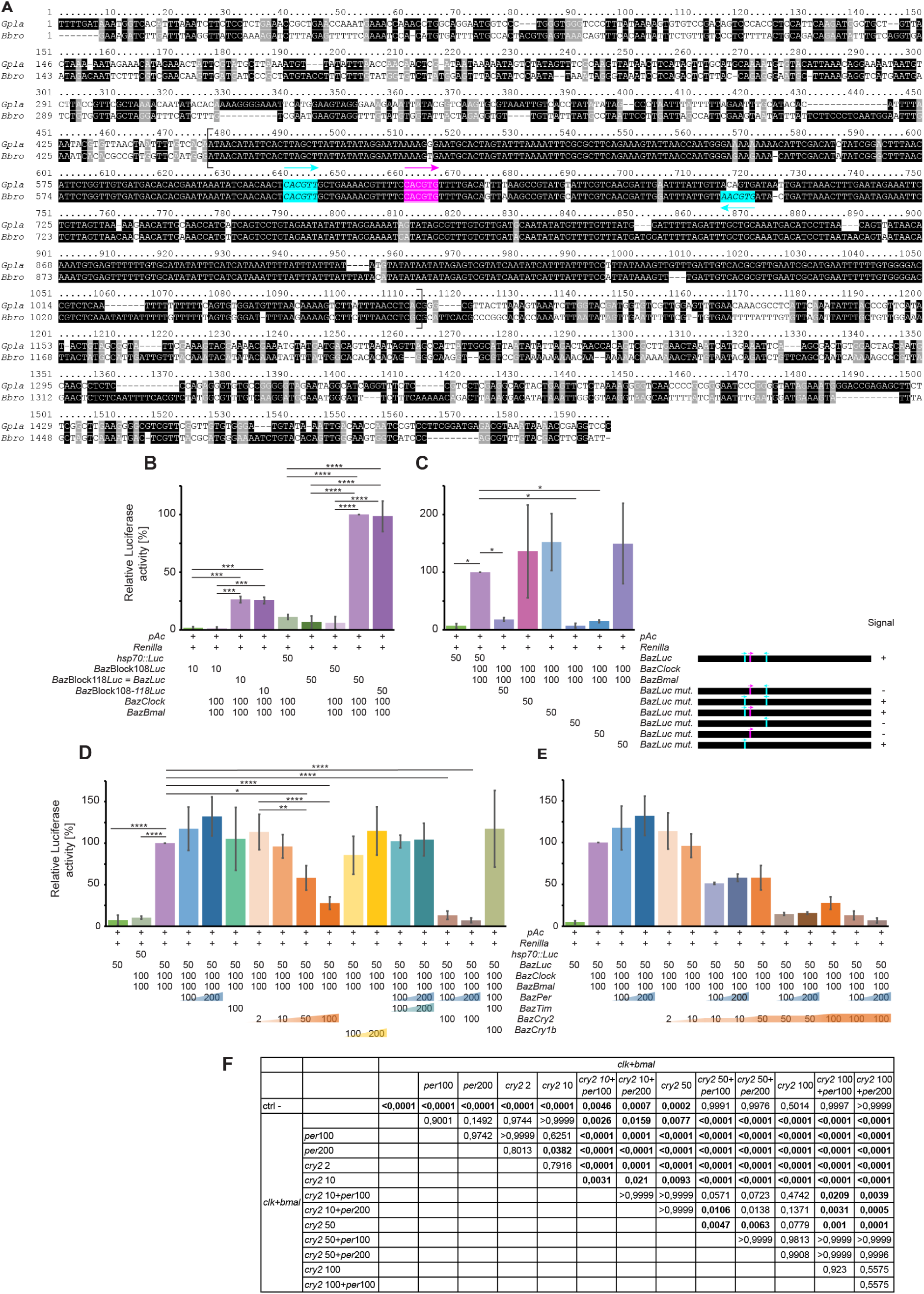
*Bathymodiolus azoricus* circadian clock genes are functional in luciferase assays using an endogenous deep-sea mussel enhancer. A. Alignments of the ortholog block 118 plus 450 nucleotides upstream and downstream in *Gigantidas platifrons* and *Bathymodiolus brooksi.* Pink underlined motif: E-box CACGTG. Turquoise italicized motifs: E-box-like CACGTT. B. *BazClock* and *BazBmal* activate the luciferase signal while co-transfected with the *Baz*Block118-*Dm-hsp70*::*Luciferase* construct (shortened to *Baz*Block118*Luc* or *BazLuc*) and the artificial *Baz*Block108-118-*Dm-hsp70*::*Luciferase* one (shortened to *Baz*Block108-118*Luc*). N = 3 transfections. *Renilla*: actin5c::Renilla. *pAc*: pAc5.1_V5-HisA. **C.** Characterization of the E-boxes-like and E-box present in *Baz*Block118-*Dm-hsp70*::*Luciferase* (*BazLuc*). The E-box-like 1 is essential for the activation by *BazClock* and *BazBmal* as the mutation of this sequence abolished the luciferase signal. Similarly, mutating this sequence in combination with the E-box or the E-box-like 2 also abolishes the luciferase signal. N = 4 transfections. **D.** Further co-transfection of *BazPer*, *BazTim*, *BazCry2*, and *BazCry1b* revealed that *BazCry2* is the main repressor. N = 3 to 4 transfections. E. *BazPer* modulates the level of repression of *BazCry2*. The conditions with *BazPer* 100 or 200 ng*, BazCry2* 2 to 100 ng, and *BazPer* 100 or 200 ng + *BazCry2* 100 ng are repeated from panel D. N = 3 to 4 transfections. **F.** Statistics for the comparisons in panel E. Statistics: one-way ANOVA with post-hoc Holm-Šídák multiple comparisons. In D, the conditions *pAc5.1_V5-HisA* and *actin5c::Renilla* have only been compared to the *BazClock* + *BazBmal* condition. Values: Mean ± SD. Plasmid amounts are given in ng. Statistics in panels B to D: 0.05: *; 0.01: **; 0.001: ***; 0.0001: ****.

*Drosophila* S2 cells do not express *clk* (*22*), *per*, nor *tim* (*58*). Hence, they can be transfected with clock gene constructs or reporters to evaluate the interactions of circadian components in a controlled, circadian TTFL-based clock-deficient cellular background (*44*). We generated three luciferase constructs: *Baz*Block108-*Dm-hsp70::Luciferase*, 2) *Baz*Block118-*Dm-hsp70::Luciferase*, and 3) *Baz*Block108-block118-*Dm-hsp70::Luciferase*. We co-transfected each of these constructs with different concentrations of *BazClock* and *BazBmal* into S2 cells (**Fig.4B**). The construct containing Block 118 resulted in a clear increase of the luciferase activity, while the construct containing Block 108 did not (**Fig.4B**). *BazClock* and *BazBmal* activated the luciferase to a similar level with the construct containing both blocks 108 + 118 (*Baz*Block108-block118-*Dm-hsp70::Luciferase*) compared to the one containing block 118 alone (*Baz*Block118-*Dm-hsp70::Luciferase,* **Fig.4B**). Also, in this setting, the amount of transfected enhancer is the limiting factor in the activation, not the amount of *BazClock* or *BazBmal* (**Fig.4B**). In order to better understand the functional relationship of the different E-box (-like) sequences, we next tested by side-directed mutagenesis which of those are functionally relevant (**Fig.4C)**. E-box-like 1 appears to be essential for the activation of the luciferase: mutating this motif alone or in combination with one of the other motifs abolishes the signal, while no difference was observed when the others were removed (**Fig.4C**). A small trend for co-operativity might exist, but this was not statistically significant (**Fig.4C).** These assays show 1) that the genome of *B. azoricus* contains a functional enhancer sequence upstream the *period* gene conserved across different species of deep-sea mussels, 2) that *BazClock* and *BazBmal* can functionally interact with this enhancer, and 3) that the first E-box-like sequence of this enhancer is crucial for its functioning in the context of circadian timing.

We then evaluated the role of the potential repressors in our system. The additional transfection of *BazCry2* markedly and significantly suppressed the luciferase activation in a dose-dependent manner (**Fig.4D**). *BazPer*, *BazTim,* and *BazCry1* alone or in combination with each other did not show any transcriptional repression/activation (**Fig.4D**). The absence of transcriptional repressive activity for *BazPer* is remarkable, given the conserved transcriptional repressor role of *period* orthologs in drosophilid flies and mammals (*3*). We thus wondered if it might instead modulate the repressive effects of *BazCry2*, which appeared as the major transcriptional repressor in this mussel. This would be particularly interesting in light of the finding that *per*, but not *cry2* transcripts (and hence likely proteins) are themselves circatidally changing under natural and lab conditions (*10*).

Our initial co-transfection of *BazPer* at 100 or 200 ng with *BazCry2* 100 ng showed no significant effect (**Fig.4D**). We wondered if this could be due to the already very high repression by *Cry2* alone and therefore tested the effect of *BazPer* together with lower concentration of *BazCry2*, 10 and 50 ng. Indeed, this revealed significant synergistic repressive activity of *BazPer* with *BazCry2* (**Fig.4E, F**). The co-transfection of *BazPer* at either 100 or 200 ng and *BazCry2* at 10 ng increased the repression of *BazCry2* alone at 10 ng. Similarly, the co-transfection of *BazPer* at either 100 or 200 ng and *BazCry2* at 50 ng increased the repression of *BazCry2* alone at 50 ng (**Fig.4E, F**). Hence, *BazPer* can modulate the repressive effects of *BazCry2*. Given *BazPer* own circatidal regulation in the lab and at the vents, this may provide a first insight into the mechanism by which a mussel’s cell autonomous circadian oscillator could switch between circadian and circatidal oscillations, depending on the relative concentrations of *BazCry2* and *BazPer* (**Fig.5**).

**Figure 5.**
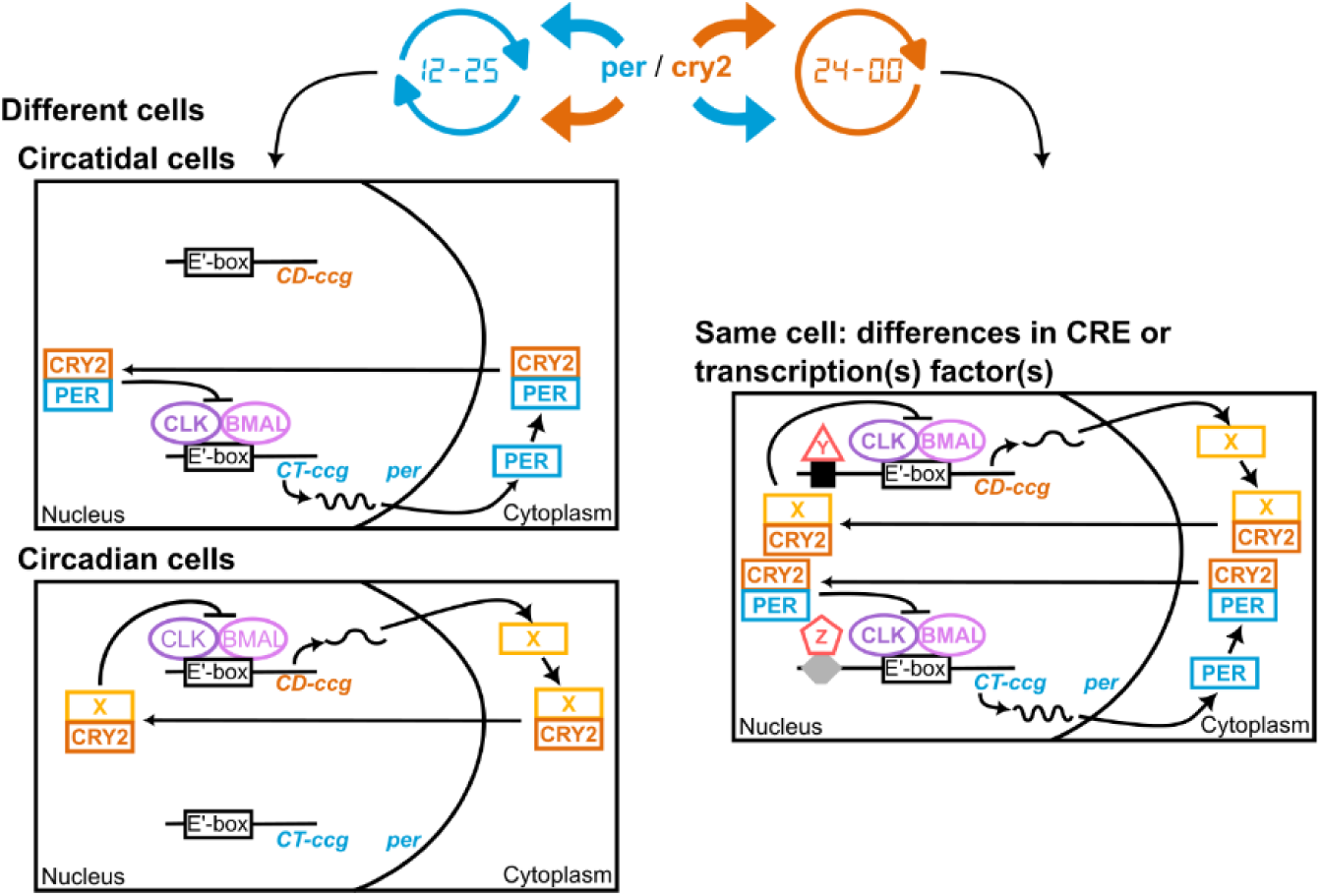
Proposed model for the generation of both circatidal and circadian rhythms in deep-sea mussel tissues. It suggests a scenario in which either different cells within a given tissue have different amounts of the respective core clock proteins, or circadian-vs. circatidal-regulated genes possess differential regulatory elements binding the respective core clock proteins with different affinities. The circatidal and circadian cells presented on the left are an extreme version of what could happen, intermediate situations could prevail. CRE: Cis-Regulatory Elements. CD: circadian. CT: circatidal. Ccg: clock-controlled genes.

## Discussion

The deeper marine organisms reside, the more extreme their environment become: while at sea level, the average pressure is 1 bar, pressure increases by 1 bar for every 10 m of depth. So, mussels living at 1700 m depth experience a pressure of 171 bars. This represents an important challenge both for *in situ* and lab experimentation, as depressurization and sea-surface pressure are stressful, often lethal, for deep-sea specimens (*59*). Consequently, most experiments on deep-sea mussels were previously performed fixing animals immediately onboard after samples recovery (*35*, *60*, *61*), or with shallower specimens for longer experiments (*62*–*66*). Here, we successfully maintained *B. azoricus* collected at −1700 m at atmospheric pressure for >1 year, an unprecedented duration for this species. Mortality happened mostly during the first month, likely reflecting an acute stress from depressurization and transport, then plateaued. This is consistent with deep-sea shrimps’ maintenance (*67*) and shows that the remaining individuals were able to physiologically adjust to the new environmental conditions. While the concept of a depth-related barrier or bottleneck is typically framed in the context of colonization from shallow to deep waters (*68*), our result challenges the commonly held view that deep-sea fauna has low tolerance to prolonged exposure at sea-surface pressure (*59*) and indicates that deep-sea fauna, or at least mussels, might have a greater pressure-tolerance plasticity than initially considered. In the lab, surviving individuals exhibited normal behaviours—active movement, regular valve-opening, foot extension. In bivalves, foot extension correlates with substrate assessment (*27*), while valve-opening correlates with respiration, nutrition, and water-quality sensing (*27*, *69*). Additionally, in the Atlantic, the Lucky Strike hydrothermal vent field is more extensively explored than the shallower Menez Gwen field, mainly because it houses a deep-seafloor observatory (https://www.emso-fr.org/EMSO-Azores), making it an ideal sampling site. Housing deep-sea mussels from 1700 m depth for up to a year in the lab is therefore a major step forward for future biological research.

The ability to successfully run temporal experiments on *B. azoricus* primary cell cultures is a key achievement, as it demonstrates the feasibility of studying circadian and other rhythms at live cell level in a deep-sea organism and may even open the door for future reverse genetic analyses. As a first step, we were able to show a circadian rhythm in transcription for foot cell cultures, an unexpected result since the strongest *in situ* evidence at vents points to tidal cycles, and the nature of any daily cue there remains unclear. Yet, the dominant signal in mussels’ cells is circadian. The proportion of identified circadian transcripts (2.8%) is actually similar to the one obtained on the deep-sea floor (2.5%; *10*). However, here it represents 157 transcripts out of 5616, while on the deep-sea floor it represented 869 out of 34 163 transcripts. Although the Smart-seq3 protocol is optimized for the detection of RNA in low RNA input or single-cells experiments (*38*), we captured only about 17% of the *B. azoricus* transcriptome. This low percentage is consistent with challenges reported for single-cell / low RNA input methods (*70*, *71*). Of note, the core circadian genes were not captured in the RNAseq data in this study. They were already lowly expressed in gill tissues (*10*), from which RNA quantities were not a limiting factor. Despite these limitations, this work shows that a circadian rhythm is endogenously present in *B. azoricus’* cells, providing complementary insights to the studies of circadian and circatidal clocks from whole organisms, like *Parhyale hawaiensis or Eurydice pulchra,* which focus on the brain as a control center (*19*). It also advances our understanding of circadian biology and its evolution in extreme environments.

Furthermore, the results from the luciferase assays show that *B. azoricus* possesses a cell autonomous functional circadian clock. In conventional model organisms like fruit flies and mice, the basic helix–loop–helix transcription heterodimer CLK-CYC or CLK-BMAL1 binds to specific DNA motifs, the canonical CACGTG E-box (*20*–*22*) or the CACGTT E-box-like or E’-box (*72*). While one E-box is sufficient for the activation of the transcription of *per* in *Drosophila melanogaster* (*20*), repeated E-box/E-box-like elements are either needed to activate mammalian *per2* (*72*) or show an additive action to enhance mammalian *per1* transcription (*73*). *Bathymodiolus azoricus* contains an enhancer sequence upstream its *period* gene, which includes 2 Ebox-like and 1 Ebox. Only the first E-box-like, but not the conventional E-box, was functionally critical. The role of this E-box-like and the other conserved sequences in the enhancer regions remain to be elucidated. Additionally, using an enhancer specific to the studied species is an original approach compared to previous work (*15*, *44*, *56*, *74*), showing that the core circadian machinery is conserved enough in deep-sea mussels to be effectively tested.

So far, *Cry2* emerged as the main repressor of *B. azoricus*’ TTFL, with a dose-dependent effect. This is similar to the mechanism described in the monarch butterfly *Danaus plexippus* (*45*) and the isopod *Eurydice pulchra* (*56*), although in *E. pulchra, per* and *tim* also act as repressors, albeit less effectively than *Cry2*. In *Platynereis dumerilii*, repression is also driven by *tr-Cry*/*Cry2* (*15*), but the roles of *per and tim* have not yet been characterized. In *B. azoricus*, the co-expression of *BazPer* and *BazTim* did not repress *BazClk* and *BazBmal* action, diverging from canonical flies models (*3*), but consistent with observations in the krill *Euphausia superba* (*74*). As in krill, *BazPer* accentuates the inhibition driven by *BazCry2* (*74*).

*BazPer* showed an about tidal expression pattern in mussels in the field (under natural tidal entrainment) and in the lab (under diel entrainment; *10*). By acting as a modulator of *Cry2*, we hypothesize that *Per* could play a role either as a common regulator between the circatidal and circadian clocks, or in the tuning of a potential single bimodal clock in deep-sea mussels. Within a single cell, the relative amounts of *BazPer* and *BazCry2* could switch the clock from a circadian to a circatidal signal or vice versa. At vents, where tides are present, the gillś transcriptome of *B. azoricus* was predominantly shaped by a tidal signal. In contrast, under LD entrainment in the lab, the number of daily transcripts was 3x higher than in the field. However, a circatidal rhythm was still present, with the number of tidal transcripts reaching 80% of the levels observed in the field (*10*). As both tidal and diel rhythms were detectable from the same tissue, we propose two possible models, depending on whether these rhythms arise in different cells or together in the same cells (**Fig. 5**). In the first scenario, there would be separate circatidal and circadian cells in the same tissue, with and without period expression. The other scenario predicts that cells can produce both circatidal or circadian rhythms, implying that the oscillation would depend on the co-binding of *per* based either on different motifs in the enhancer or different transcription factors. It could also be a combination of both possibilities.

Several studies increasingly support the idea that canonical circadian core clock components may not be intrinsically circadian. Instead, their oscillations may depend on the organism’s dominant cycle (*75*–*77*). This thus includes *BazPer* (*10*). The mechanism of the circatidal clock remains unidentified. Three hypotheses have been proposed to explain its functioning. Naylor proposed the existence of 2 distinct biological clocks, a circatidal clock and a circadian one (*78*). Palmer and Williams proposed the co-existence of two circalunidian (∼24.8h) clocks working in antiphase to generate 12.4h tidal rhythms (*79*, *80*). And Enright suggested that a single, flexible circadian clock with a bimodal nature could generate both circadian and circatidal rhythms (*81*, *82*). Experimental manipulations in crustaceans and insects support the existence of distinct circadian and circatidal clocks. In the mangrove cricket *Apteronemobius asahinai* or the isopod *E. pulchra*, disrupting core circadian components abolishes circadian rhythms while leaving circatidal patterns intact (*56*, *83*, *84*). Similarly, environmental cues such as light or vibration can separate circatidal and circadian cycles (*56*, *57*). Yet the picture is complicated by shared molecular players—casein kinase 1 inhibition impairs both rhythms in *E. pulchra* (*56*), and BMAL1 is required for the circatidal rhythm of both *P. hawaiensis* (*57*) and *E. pulchra* (*85*). The latest work on *E. pulchra* and *P. hawaiensis* even implies a stronger intertwining of both systems: separate groups of cells in the brain of these crustaceans can show either a circadian rhythm of clock gene expression, or a circatidal rhythm of expression for *Eptim* or *PhPer* and *PhCry2*, suggesting separate circadian and circatidal neural clockworks (*19*). We propose that Enright’s hypothesis of a single clock functioning in deep-sea mussels, where the *Cry2/Per* ratio could induce either a circadian or a circatidal oscillation, is at present most consistent with our data. However, both models might very well be valid. Mussels possess a quite simple nervous system that relies on 3 pairs of ganglia without a centralized brain such as in crustaceans (*27*). Hence, the circatidal and circadian clock mechanisms may vary in different taxa. In the oyster *Crassostrea gigas*, it has already been proposed that Enright’s hypothesis might prevail, and that circatidal rhythms may arise from a circadian clock synchronized to tidal cycles (*77*, *86*). A similar pattern emerges in shallow-water mussels: only a weak circadian rhythm has been shown in valve-behaviour in *Mytilus edulis* (*87*), while no circatidal clock has yet been demonstrated. Field and laboratory studies of shallow-water mussels describe that transcript abundance, gill metabolism, cardiac activity, and protein levels all oscillate predominantly on a 24-hour cycle, even when tidal fluctuations are superimposed (*88*–*93*). Future analyses, involving genome analyses and core “circadian” protein localizations in mussels’ cells and tissues, will allow to discriminate the different scenarios.

## Conclusion

We show that tissue cultures from deep-sea mussels exhibit significant cell autonomous circadian transcript cycles, a surprising finding given that tidal, not diel, cycles are detectable in their natural habitat. Based on experiments that tested a *B.azoricus* endogenous enhancer region with core circadian clock genes in a heterologous tissue culture assay, we suggest that *BazPer* can modulate *BazCry2*, the latter acting as the major repressor in the deep-sea mussels’ cellular circadian clock. If Cry2 is present at high concentrations, Per’s impact will be negligible. Yet, at lower Cry2 concentrations, Per may significantly impact on the core clock loop dynamics, which is particularly interesting given that *per* transcripts show themselves ∼tidal rhythms under natural deep-sea conditions and in the lab. This may provide a first idea of how a circadian clock could switch between different periods, even in a tissue autonomous, nervous-system independent manner. Overall, our work on a deep-sea ecological key species opens new venues to explore how life can adapt to the most extreme conditions, in which the right timing makes the difference between life and death.

## Acknowledgments

The authors acknowledge the captain and crew of the RV Pourquoi Pas? and RV L’Atalante, the pilots of the HOV Nautile and ROV Victor6000 of the Momarsat 2022 and 2023 cruises, and Jozée Sarrazin for their sampling contribution. They also thank the NGS Team from the Vienna Biocenter Facility for the RNA sequencing, Alexander W. Stockinger and Petra Schaffer for the adaptation of the dissociation protocol for cell cultures, Niklas Fudikar for his help cloning the core clock genes of *Bathymodiolus azoricus*, Julie Tourolle for providing the map in Fig.1, and Michael Bindl, Lukas Orel, Pedro Bum, and Irmgard Fischer for their input on cell cultures. Franziska Lorbeer and Alexander Stark kindly provided the *hsp70* core promoter of *Drosophila melanogaster*.

## Funding

VBC post-doctoral program VIP2 co-funded by the Horizon 2020 Marie Skłodowska-Curie Grant Agreement #847548 (AMM).

Helmholtz Society, distinguished professorship by the Alfred Wegener Institute Helmholtz Centre for Polar and Marine Research (KT-R).

H2020 European Research Council, ERC Grant Agreement #819952 (KT-R). Austrian Science Funds (FWF), SFB F78 (KT-R).

Human Frontier Science Program (HFSP), #RGP021/2024, https://doi.org/10.52044/HFSP.RGP0212024.pc.gr.194174 (KT-R, MM).

The Momarsat cruises are supported by the French Oceanographic Fleet.

None of the funding bodies was involved in the design of the study, the collection, analysis, and interpretation of data or in writing the manuscript.

## Authors contribution

Project design and funding acquisition: AMM, MM, KTR. Mussels sampling: MM. Experimental design: AMM, KTR. Establishing cell cultures: AMM. Temporal experiment: AMM, FS. RNA extraction: AMM. Bioinformatic analyses: CK. Biostatistics and data analyses: AMM. Luciferase assays: AMM, FS. Data interpretation: AMM, FS, KTR. Manuscript original draft: AMM. Manuscript review and editing: AMM, KTR, FS, MM, CK. All authors read and approved the final manuscript.

## Data and materials availability

The full CDS and the enhancer sequences have been submitted to NCBI (accession): *Clock* (PV892275), *bmal* (PV892276), *period* (PV892277), *timeless* (PV892278), *cryptochrome1b* (PV892279), *Cryptochrome2* (PV892280), Block 108 (PV892281), Block 118 (PV892282). The RNAseq data are available at NCBI/SRA under the BioProject accession PRJNA1308002.

## Supplementary Information

**Figure S1.**
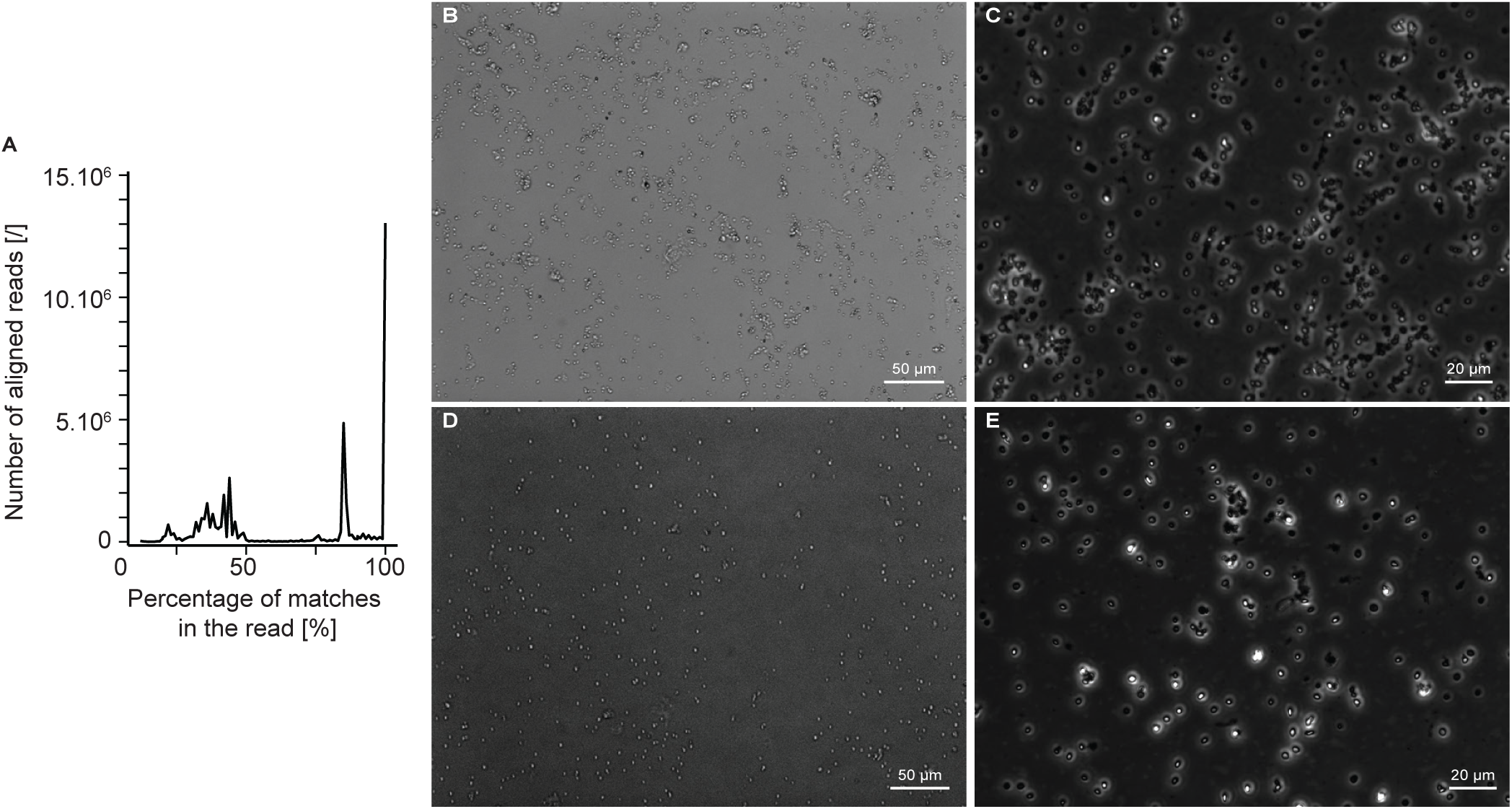
Quality control of several the foot cultures. **A.** Control of the culture after RNA sequencing. The X axis represents the percentage of matches in the reads from 0 to 100 %, calculated as the number of matched nucleotides divided by the total length of the read (150 nt). The Y axis represents the number of aligned reads for each matched fraction. **B, C** and **D, E.** Images of two distinct foot cultures imaged after 951 days in the lab.

**Figure S2.**
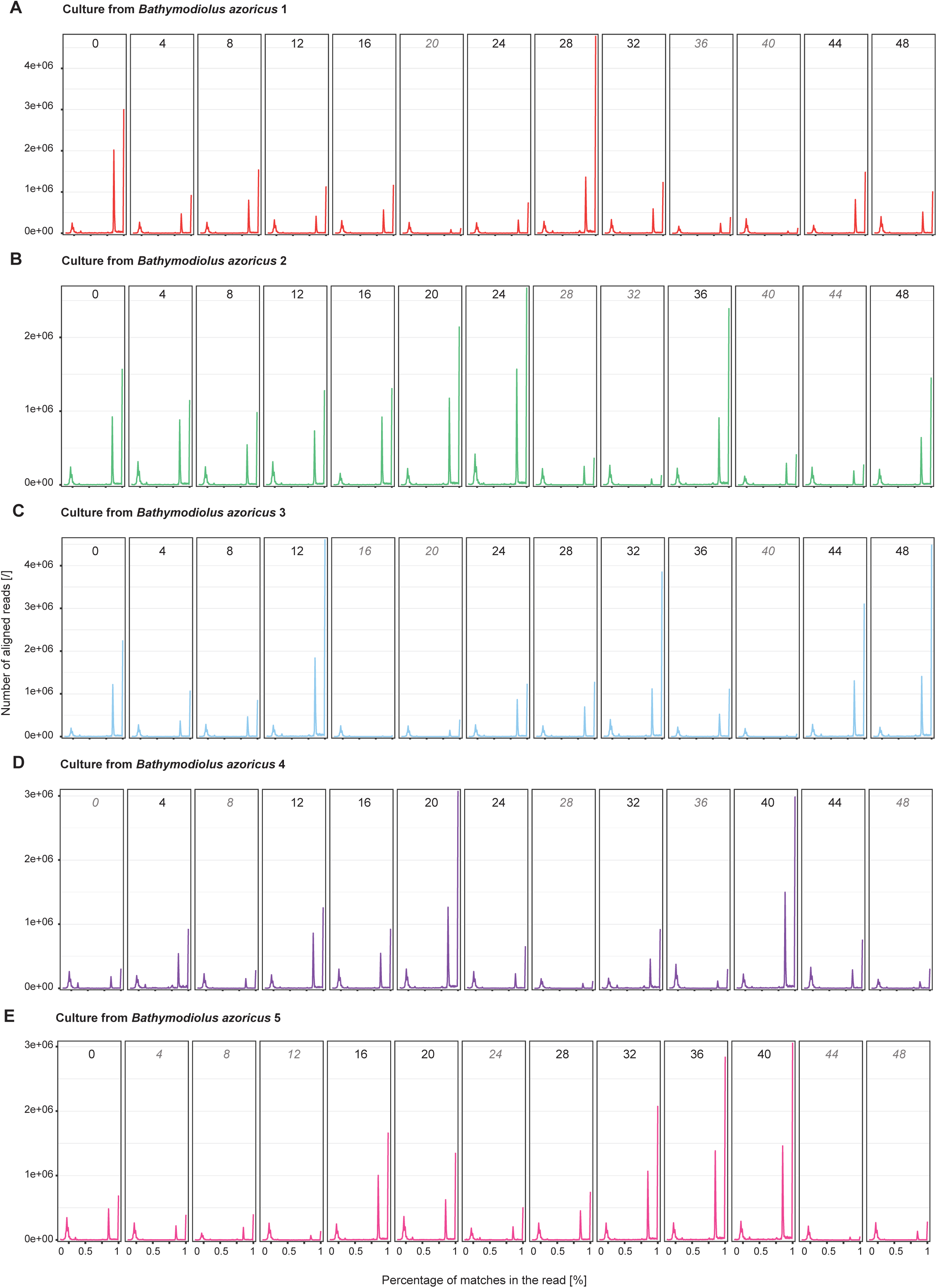
Quality control of the RNAseq samples used for the temporal cell culture experiment. Foot tissues from 5 different mussels were placed in culture in constant darkness at 4°C, sampled over 48h after 25 days, and RNA was extracted and sequenced. For each culture, the X axis represents the matched fraction from 0 to 100 %, calculated as the number of matched nucleotides divided by the total length of the read (150 nt). The Y axis represents the number of aligned reads for each matched fraction. Italics: samples with less than 500k alignment counts at the 150 nt peak and that were therefore filtered out.

**Figure S3.**
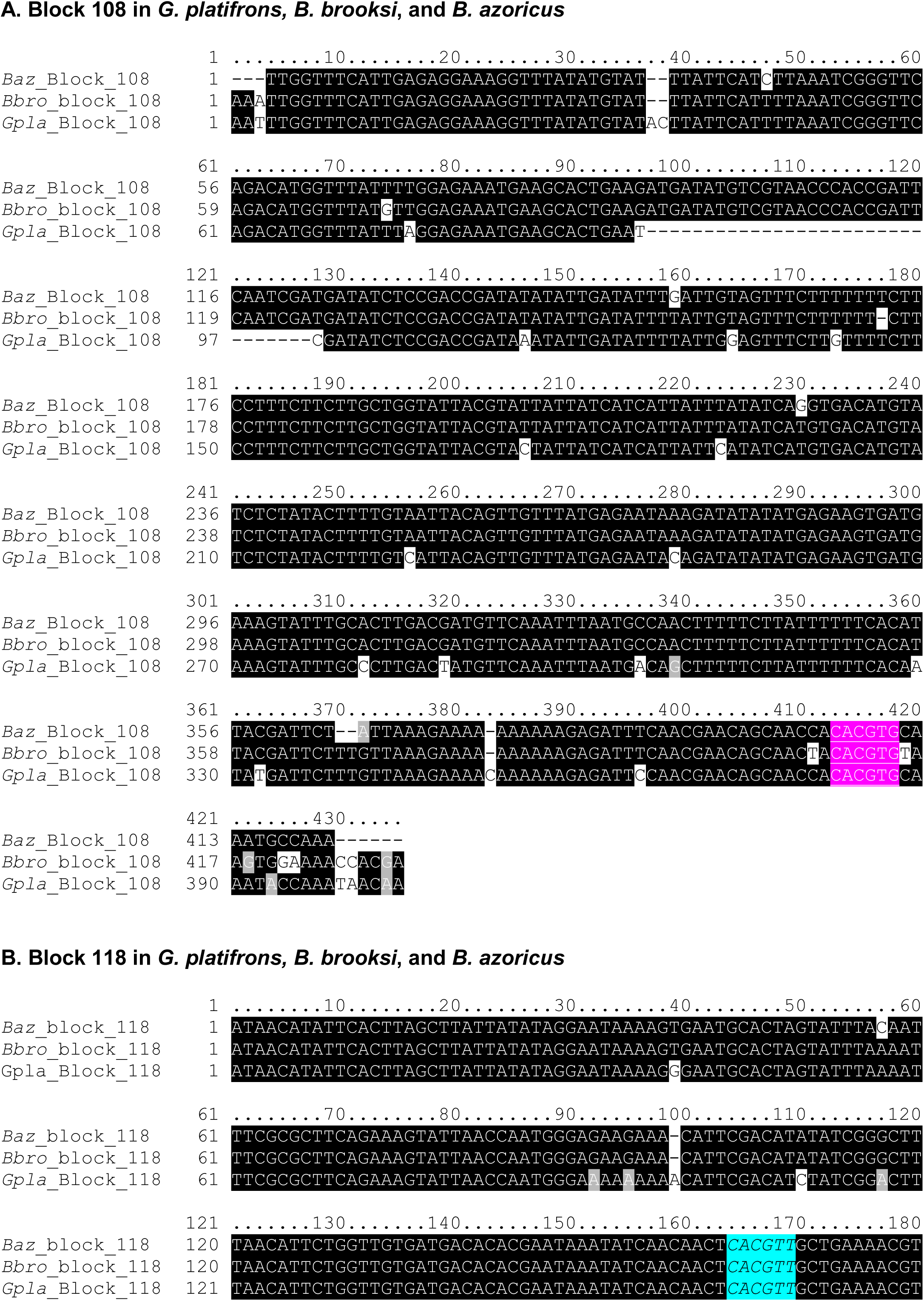
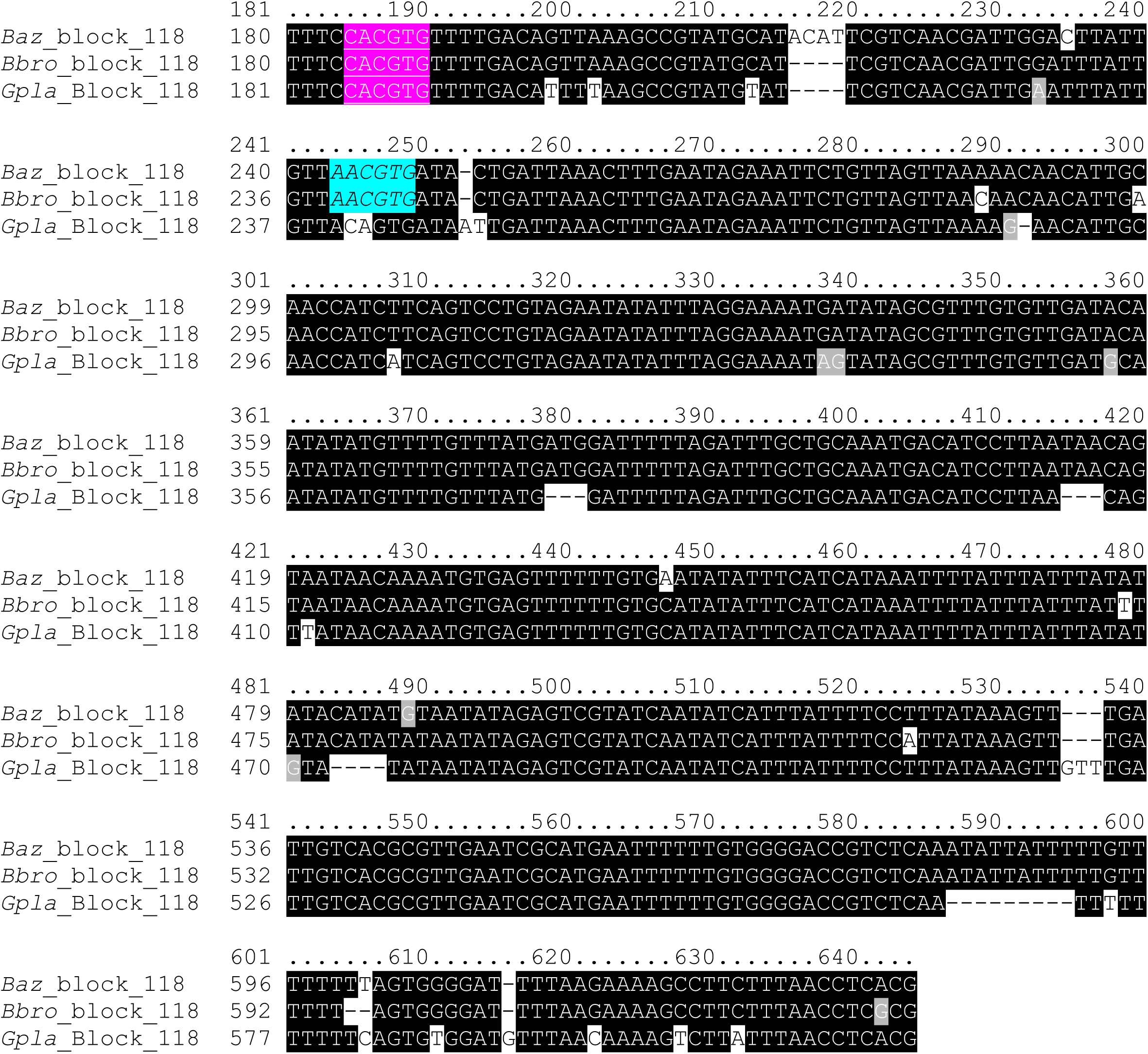
Alignments of the 2 ortholog blocks containing a CACGTG E-box in *Gigantidas platifrons, Bathymodiolus brooksi,* and *Bathymodiolus azoricus.* Pink underlined motif: E-box CACGTG. Turquoise italicized motifs: E-box-like CACGTT.

---

**Table S6.** Comparison of the results of the rhythmic analysis on the actual dataset to the results of the rhythmic analysis after shuffling the columns of the dataset. The procedure has been bootstrapped 10 000 times. CD: total number of circadian transcripts. CT: total number of circatidal transcripts. CD-R or CT-R: transcripts that are now identified as circadian or circatidal and that were already identified as rhythmic (either circadian or circatidal) in gill RNAseq data on the deep seafloor or in the lab (Mat et al, 2020). CD-CD: transcripts that are now identified as circadian and that were already identified as circadian in gill RNAseq data (Mat et al, 2020). CT-CT: transcripts that are now identified as circatidal and that were already identified as circatidal in gill RNAseq data (Mat et al, 2020).

| **Circadian analysis** |  |  |
| --- | --- | --- |
| **Parameters** | **Actual values** | **Actual > shuffled & bootstrapped** |
| CD | 157 | 70.3 % |
| CD-R | 66 | 77.9 % |
| CD-CD | 40 | 90.8 % |
| CD and CD-R | 157; 66 | 81.4 % |
| CD and CD-CD | 157; 40 | 92.1 % |
| CD and same CD-R | 157; same 66 | 100 % |
| **Circatidal analysis** |  |  |
| **Parameters** | **Actual values** | **Actual > shuffled & bootstrapped** |
| CT | 109 | 6.8 % |
| CT-R | 41 | 8.5 % |
| CT-CT | 26 | 31.3 |
| CT and CT-R | 109; 41 | 11.4 % |
| CT and CT-CT | 109; 26 | 32.4 % |
| CT and same CT-R | 109; same 41 | 100 % |

